# An Array of Gut-on-a-Chips for Drug Development

**DOI:** 10.1101/273847

**Authors:** Nureddin Ashammakhi, Mohammed Elmusrati

## Abstract

Drug development is a time and cost-consuming long process. Conventionally, this involves the use of two-dimensional (2D) cell culture and animal testing. However, these methods often fail to mimic normal human physiology and drug metabolism, and sometimes complications may occur later because they do not appear in preclinical studies. With recent development of organ-on-a-chip devices, these biomimetic systems try to recapitulate human physiology more than other methods, and promise to largely replace animal studies and reduce costs and complications. Because gut is an important route for drug administration and absorption, gut-on-a-chip platform was developed. To develop a high throughput system, a recent work reported on using a multiplication array of several chip devices, demonstrating the efficiency in studying drug effect on gut barrier function as compared to conventional methods. Huge data was generated, and thus new multidisciplinary approach is needed where data collection, communication and analysis platforms need to be developed. This will enable validation of organ-on-a-chip approach and lay basis for its establishment as efficient and cost-effective tool in drug development. Because of the importance of the work, this commentary highlights important aspects of the topic and try to stimulate new the development of innovations in the field.

**Disclosure:** No conflict of interest.

## 1. Introduction

Current methods for assessing toxicity rely on the use of 2D culture and on the use of animal experiments. It is a long and costly process to get single medicine to the market. With new EU regulation (REACH policy for Registration, Evaluation, and Authorization of Chemicals) requesting the testing for toxicity of ∼30,000 chemicals, we will need about 2.5-54 million animals ^1^ to use for testing [costing about 1.3-9.3 trillion Euros ^2, 3^]. Animal studies are costly, slow, associated with ethical questions and fail to predict human responses. Using animals is limited also by species-dependent variations. It is also difficult to extrapolate results obtained from animal studies to humans ^4^. Unfortunately, many drugs that may proof safe in conventional cell culture studies and in animal studies, result in side effects appearing later during clinical trials or even after their approval and use. Thus, using a more biomimetic testing method such as tissue or organ on chip platform is appealing. In new US regulations, e.g. Toxicology in the 21st Century (Tox21) program calls to move away from animal studies^5^.

## 2. Gut-on-a-chip

With the recent advances made in microfluidics ^6^ and their application in developing lab-on-a-chip ^7^ which was then combined with cells to have “organ-on-a-chip” ^8^ devices, it is becoming evident that such devices can be used as an alternative to animal experiments and provide large volume of data and can possibly be more biomimetic that animal experiments themselves as they employ human-derived cells.

Because many drugs are absorbed through the gut, gut-on-a-chip devices developed in 2012-2013 ^9^,^10^ that has also flora included ^11^ to create such a testing platform. The system is comprised of intestinal epithelial cells cultured in one side of a membrane in the middle of a microchannel contained in a microchip. Fluid flows on both side to represent luminal and basal sides of the epithelial lining of the gut (Figure 1a & b). This achievement represents important an important milestone that following the idea developed by the same group in 2010 for lung-on-a-chip ^8^.

**Figure 1.**
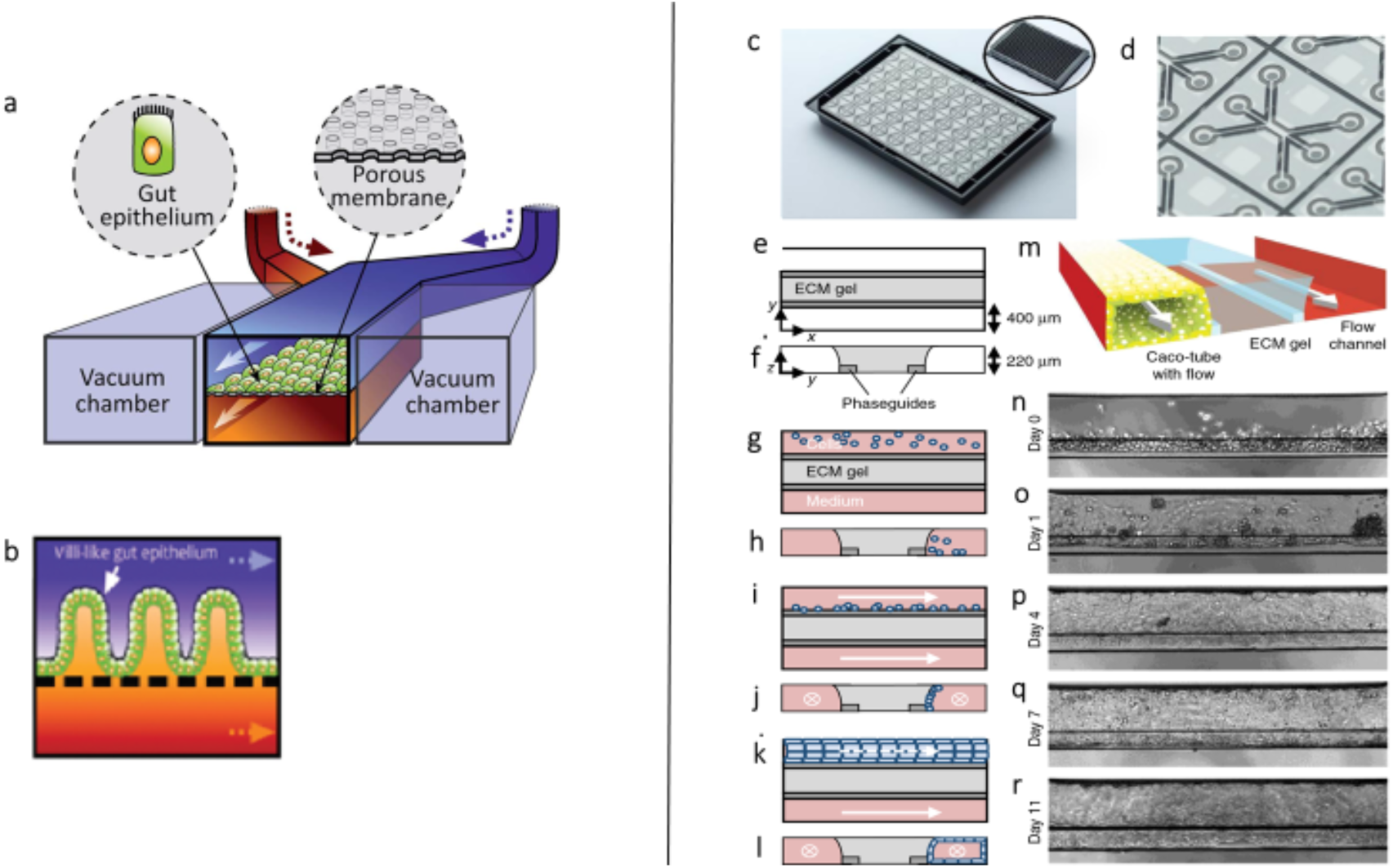
a) Schematic illustration of the gut-on-a-chip device which comprises several channels. The central channel has a flexible porous ECM-coated membrane. Gut epithelial cells were cultured on the top of the membrane. Vacuum chambers are seen laying on both sides of the central channel. Reproduced with kind permission form Kim *et al.* ^11^, b) Schematic illustration showing the formation of intestinal villus structure. Reproduced with kind permission form Kim *et al.* ^10^ c) OrganoPlate system having 40 microfluidic channel networks. The inlay shows top view of the 384-well plate format device. d) Single microfluidic network having three channels joined in the center. e, g, i, k) Horizontal projection and f, h, j, l vertical cross section of central region for the following steps. e, f) An extracellular matrix (ECM) gel (light gray) is patterned by two phaseguides (dark gray). g, h) Lanes adjacent to the ECM gel had culture medium introduced into them. One of the lanes has cells. i, j) Cells were allowed to settle against the ECM gel by placing the plate on its side. k, l) Following flow, cells form a confluent layer that lined the surface of channel and gel. m) Schematic of the chip center comprising a tubule, ECM gel and a perfusion lane. Two phaseguides (white bars) define the three distinct lanes in the central channel. At its apical side, the tubule has a perfusable lumen. n–r) Images (phase-contrast) of the tubular structure formation at day 0, 1, 4, 7, and 11, respectively. Scale bars: 100 μm. Reproduced with kind permission form Trietsch *et al*. (1).

In addition to organ-on-a-chip models mimicking normal physiology, organon-a-chip models for pathological conditions such as inflammation ^12^, thrombosis ^13^, degeneration ^14^, and cancer ^15^ can also be developed and used for drug development studies.

## 3. Multiplied organ-on-a-chip devices

In next important logical step, which also constitutes the second milestone, recently Trietsch *et al*. reported on a new device comprising an array of gut-on-a-chip joined together (357 gut tubes in 10 plates) and the efficiency of this setup in testing for drug toxicity was demonstrated ^16^.

It is for the first time that such comprehensive method is used for assessing drug concentration and exposure time on the integrity of intestinal epithelial barrier. It is the largest ever reported organ-on-a-chip system that used more than 20,000 data-points. The study opens new avenues for efficient and reliable future drug testing and development. Trietsch *et al*. ^16^ have cultured human intestinal colorectal adenocarcinoma cell line (Caco-2) in one microfluidic channel and were separated by collagen-type-I gel from fluid flow channel (Figure 1c-r). Ten plates, each comprising 40 leak-tight tubes in parallel, were used. Cells exhibited differentiation characteristics and markers such as brush border, tight junctions, fluid transport and intact epithelial barrier function. Cell polarization against the gel was also observed. Fluorescent probes were used to assess barrier integrity and its drug-induced loss of integrity with resulting leak at day 4 of culture. Barrier integrity loss was assessed by exposing the apical side of cell lining to Staurosporine (inducer of apoptosis) and aspirin (affects tight junctions) for 125 h. Comparison experiments were performed in conventional Transwell and showed lower sensitivity to detect barrier disruption as compared to organ-on-a-chip setup. A clear drug concentration-response effect was also detected. Although such organ-on-a-chip devices offer a new platform for drug testing, the technology needs validation against current standards and clinical results to establish a solid recognized method of testing in drug development. This work ^16^ is thus also important contribution in this direction.

## 4. Data Analysis

While the current work multiplies the organ-on-a-chip units many times creating huge and unprecedented amount of data, it requires parallel development of appropriate data analysis and management system as well as modelling. New multidisciplinary approach is thus needed and should be based on developing at least the following layers: 1. Sensing technologies, to sense and measure the cell responses accurately; 2. Communication technology, to collect time series data from maybe tens of thousands of sensors effectively; 3. Big data analysis and mining using effective artificial intelligence and machine learning algorithms; 4. Biochemical analysis and modelling, for drug design, testing and evaluation; And, 5. Pathological models for the assessment of changes and effect of drug(s) in pathological situation(s). There is a different optimisation and working procedures of each layer. However, cross-layer optimisation would be crucial to enhance the efficiency and the speed of the whole process. Another possible important application of the system is the online medical drug design or fine-tuning. The testing phase will just give us a conclusion about the validity of the drug and its effectiveness as well as possible side effects on the short and long-terms. However, with appropriately large number of data-points with many related sensors involved and online data analysis (using for example advanced machine learning and intelligent automation), it could be possible to modify the drug (e.g., chemical structure, compound concentration, design process) online. Within the closed-loop intelligent operations of drug-modifications and their impacts on cells, we will be able to perform thousands of directed (optimised) trials within proper duration of time. Current technical systems in terms of reliable electronics, sensors accuracy, computing facilities, smart algorithms, and intelligent micro-automated systems can be integrated as the foundation of the next generation drug design and development.

## 5. Conclusions

Gut on chip was developed to assess drug testing and aid drug development. To achieve high throughput, multiplications array of these devices were recently developed and its efficiency in such process was demonstrated. Data generated is huge and needs development of proper management. Multidisciplinary approach is necessary to develop novel functional platforms for drug testing and development based in multiplied organ-on-chips, data collection, communication and analysis. Costs will hence, be saved, efficient medicines developed and risks to patients tremendously reduced.

